# Characterization of a GH17 laminarinase, *Ml*GH17A, from a laminarin polysaccharide utilization locus in the marine bacterium *Muricauda lutaonensis*

**DOI:** 10.64898/2026.01.22.701063

**Authors:** Leila Allahgholi, Maik G.N. Derks, Antoine Moenaert, Zehui Dong, Justyna M. Dobruchowska, Javier A. Linares-Pastén, Ólafur H. Friðjónsson, Guðmundur Óli Hreggviðsson, Eva Nordberg Karlsson

**Author notes:** For correspondence: Eva Nordberg Karlsson, or.

## Abstract

Heterotrophic marine microorganisms have the capability to degrade and metabolize laminarin, which is the most abundant source of energy and nutrients in the marine environment, via enzymes encoded by genes clustered in polysaccharide utilization loci (PULs). In this study, a PUL potentially responsible for laminarin utilization was identified in the genome of the marine bacterium *Muricauda lutaonensis* strain ISCAR-4703, with conserved synteny in the genus *Muricauda.* A GH17 laminarinase (*Ml*GH17A) encoded in the newly identified PUL was cloned, produced, and characterized as an endo-acting laminarinase, exhibiting the ability to degrade laminarin and laminari-oligosaccharides with a degree of polymerization (DP) greater than four into laminaribiose, laminaritriose, and laminaritetraose, with laminaritriose as the main product making up >50% of the produced oligosaccharide products. The three-dimensional model of the enzyme revealed the presence of seven putative subsites, including four glycone subsites (-4 to -1) and three aglycone subsites (+1 to +3), with a wide cleft to accommodate branches at the -2 subsite, enabling it to act on β-1,3 linked backbones in polysaccharides with β-1,6 linked branches. This enzyme is, along with the recently characterized β-1,3 glucanosyltransglycosylase (*Ml*GH17B), conserved in several *Muricauda* species and is suggested to play a crucial role in the utilization of laminarin by these bacteria.

**Importance:** Laminarin, a β-1,3-glucan with occasional β-1,6 branching, is the most abundant source of energy and nutrients in the marine environment. In this study, polysaccharide utilization loci (PULs) for laminarin degradation were identified in various marine *Muricauda* species, encoding a range of glycoside hydrolases and transglycosylases. In *Muricauda lutaonensis* ISCAR-4703, the PUL included two GH17 enzymes, separated by a GH30 enzyme and a major facilitator superfamily (MFS) transporter, a feature observed in all corresponding *Muricauda* PULs. A novel endoacting laminarinase from the PUL, *Ml*GH17A, was characterized and shown to hydrolyze laminarin into laminaribiose, laminaritriose, and laminaritetraose, with laminaritriose as main product. Bioinformatic analysis showed that the enzyme lacked the typical subdomain found in GH17 plant β-glucanases, leading to a lower number of aglycone subsites (+1 to +3). Instead, *Ml*GH17A possessed more glycone subsites (-1 to -4), attributed to the β3-α3 loop, which was longer than in GH17 plant β-glucanases.

## Introduction

Laminarin is a carbohydrate polymer, with a degree of polymerization (DP) of approximately 20–35 glucose units, primarily found in brown algae (Phaeophyceae) and certain species of marine diatoms that serves as a storage carbohydrate and energy reserve in these organisms (1). Laminarin, considered a dietary fiber, is composed of a β-1,3-D-glucan backbone with sporadic β-1,6-linkages and reducing ends capped with either a mannitol or glucose moiety (2).

Laminarin and laminari-oligosaccharides are reported to exhibit several bioactive properties, such as prebiotics, anti-tumor, anticancer, anti-inflammatory, and antioxidant properties (3). However, these characteristics are contingent upon their degree of polymerization and the level of branching (4). Laminarin-utilizing enzymes such as laminarinases or β-1,3-glucanases and β-1,3-glucanosyltransglycosylases found in microorganisms, including bacteria, archaea, and fungi, present a viable opportunity for the production and modification of laminari-oligosaccharides with potential bioactive properties.

Laminarin is the major molecule in the marine carbon cycle (2). The ability to utilize laminarin is vital for microorganisms that live in aquatic environments where laminarin is abundant. It allows them to efficiently obtain energy and nutrients from their surroundings, helping them to survive and thrive in these environments and aiding in the prevention of the build-up of excess organic matter in the marine environment (2, 5). This is important because excessive organic matter without balanced bacterial or enzymatic degradation can lead to ecological imbalance and in certain ecological niches, result in harmful algal blooms and oxygen depletion (6). Microbes have specific pathways for utilizing polysaccharides, composed of enzymes responsible for breaking down various glycosidic linkages. These enzymes are often encoded by genes clustered in operons called polysaccharide utilization loci (PULs). A PUL is a strictly regulated, colocalized gene cluster that encodes enzyme and protein ensembles required for the saccharification of complex carbohydrates (7). Recent studies indicate that each type of polysaccharide requires its own corresponding PUL to be effectively degraded by a microbe. It is, however, worth mentioning that polysaccharide degrading enzymes may also be scattered across the genome and are not always located within PULs. The microbial genome must contain approximately one gene for each enzyme that targets a unique glycosidic bond in the glycan structure. This results in a direct correlation between the complexity of the targeted glycans and the number of enzyme encoding genes (8). Laminarinases, encoded in a PUL for laminarin degradation, have been identified in a bacterium from the genus *Formosa* (1) and encode enzymes that hydrolyze the β-1,3 glycosidic bonds present in laminarin, allowing the microorganism to use it as a source of energy and carbon. However, there is a lack of information about the PULs responsible for laminarin degradation in most other marine species, such as species belonging to the genus *Muricauda*, despite the isolation of numerous *Muricauda* species from diverse marine environmental conditions, including saline environments such as intertidal or tidal sediment and sand, salt lakes, crude oil-contaminated seawater, coastal hot springs, surface seawater, surface marine snow, Antarctic seawater, sponges, and the rhizosphere of marine macroalgae (9).

Among *Muricauda* species, our interest in *Muricauda lutaonensis* has risen, as it was discovered that the genome of the *M. lutaonensis* strain ISCAR-4703, isolated from a geothermal intertidal region in Iceland, encodes a GH17 β-1,3-glucanosyltransglycosylase (*Ml*GH17B), capable of producing β-1,6 branching or kinking on β-1,3-oligosaccharides (10). The current work shows that this enzyme is associated with a laminarin PUL in *M. lutaonensis* ISCAR-4703. Furthermore, in a previous genome sequencing project (11), it was noted that *M. lutaonensis* strain KCTC22339, isolated off the coast of Taiwan, contained a gene encoding a laminarinase-like protein homolog, suggested to potentially be capable of degrading β-1,3-glucans. In the current study, the genome of *M. lutaonensis* ISCAR-4703 was thoroughly investigated to determine the presence of any PUL responsible for laminarin degradation and any potential laminarinase-like enzymes for laminarin degradation and/or modification of oligosaccharides, revealing similarities to a laminarin utilization PUL of *Formosa* sp. Hel1_33_131 (1). Ultimately, a novel GH17 laminarinase from the laminarin PUL of *M. lutaonensis* ISCAR-4703 was characterized with respect to its product profile and potential role in the PUL.

## Results

### Polysaccharide utilization loci for laminarin degradation in *Muricauda* species

The genome of *Muricauda lutaonensis* (strain ISCAR-4703) was annotated to identify potential gene clusters and/or polysaccharide utilization loci (PULs). As a result, a specific PUL was discovered, as specified in Fig. 1, that includes a cluster of genes encoding enzymes relevant to laminarin utilization. Two enzymes from the GH17 family were encoded in the PUL: *Ml*GH17A and *Ml*GH17B. The enzyme *Ml*GH17B was recently shown to be a β-1,3-glucanosyltransglycosylase (10), motivating further analysis of its GH17 homolog (*Ml*GH17A) in the PUL to clarify their respective role.

**FIG 1.**
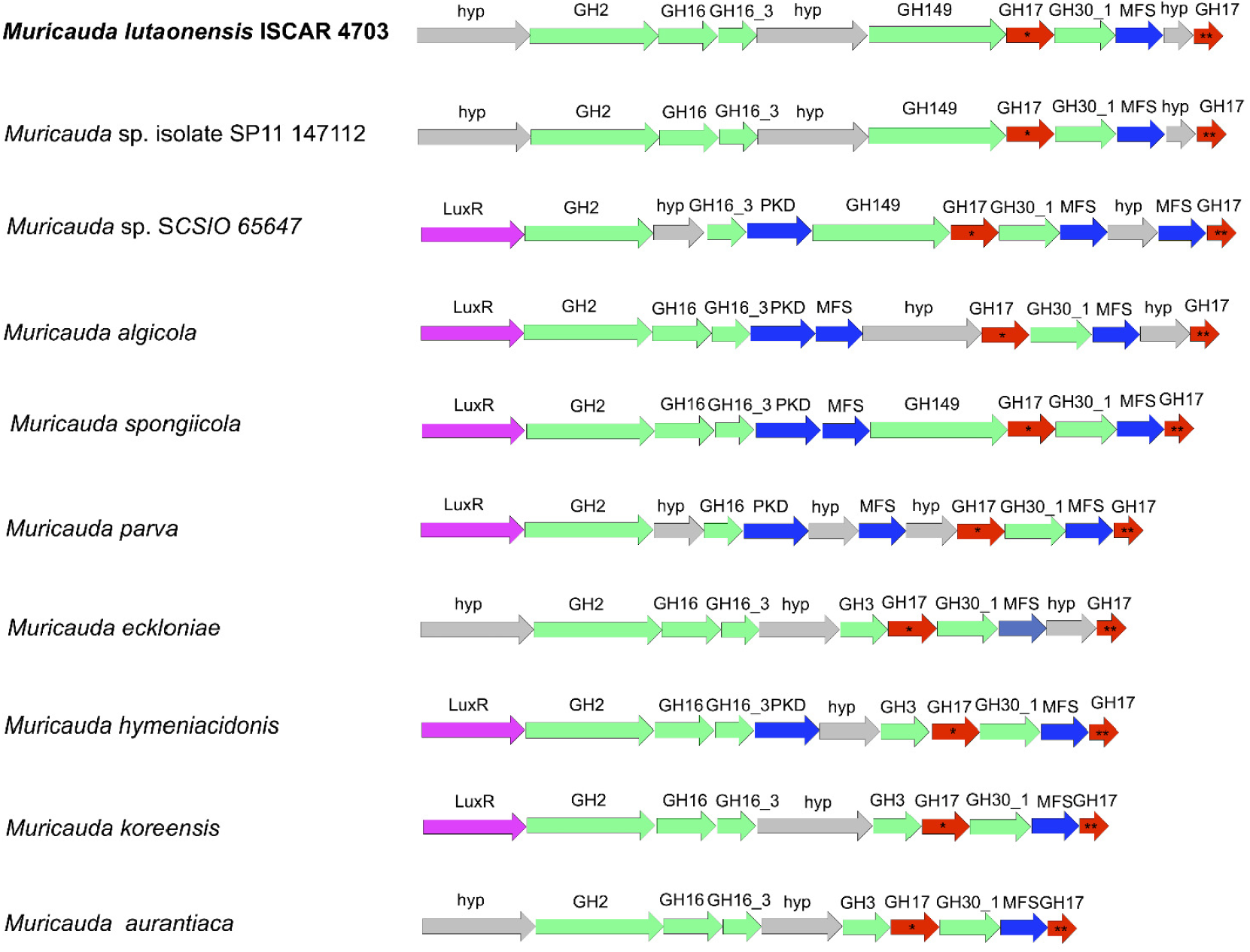
Gene clusters available in *Muricauda* species/strains. MlGH17A and *Ml*GH17B and their homologues are shown by (*) and (**) respectively.

A similarity search in the NCBI BLAST suite (12) by blastp, using the *Ml*GH17A and *Ml*GH17B sequences as query sequences against the non-redundant protein sequence database, revealed that genes coding for corresponding enzymes were conserved in other species in the genus *Muricauda,* for example *Muricauda spongiicola* strain 2012CJ35-5 (NCBI accession No. JAMFMA010000001.1), *Muricauda algicola* strain AsT0115 (NCBI accession No. VCNI01000001.1), *Muricauda eckloniae* strain DOKDO 007 (NCBI accession No. LCTZ01000001.1), *Muricauda parva* strain DSM 25885 (NCBI accession No. NZ_OBEH01000001.1), *Muricauda aurantiaca* strain HME9304 (NCBI accession No. NZ_CP030104.1), *Muricauda koreensis* strain ECD12 (NCBI accession No. QGEG01000001.1), *Muricauda hymeniacidonis* strain 176CP4-71 (NCBI accession No. SGIU01000001.1), *Muricauda* sp. SCSIO 65647 (NCBI accession No. NZ_CP091037.1), and *Muricauda* sp. isolate SP11 147112 (NCBI accession No. PARJ01000010.1) (Fig. 1).

Further analysis of the genome of the respective microorganism showed that the gene clusters displayed a high level of synteny, giving a further indication of a common role. Moreover, the position of the genes in the *M. lutaonensis* PUL largely corresponded to those in a cluster identified in a *Formosa* strain from the *flavobacteriaceae* family, where expression of the genes was shown to be induced in the presence of laminarin (1). This suggests that the PUL plays a crucial role in the degradation of laminarin by *M. lutaonensis*, and related marine bacteria.

The genes coding for the homologues of *Ml*GH17A and *Ml*GH17B in *M. lutaonensis* were always located in the downstream part of the cluster, being separated by genes encoding a GH30 subfamily 1 enzyme (GH30_1), and a major facilitator superfamily (MFS) transporter (commonly identified as a low DP sugar/oligosaccharide transporter protein (13)), respectively. Candidates from GH30_1 have, according to CAZy-classification, unambiguously been characterized as β-1,6-glucosidases (14–16). These four genes, together, constitute the main downstream part of the PUL, and the arrangement of the genes is conserved in all *Muricauda* strains. Upstream of the described genes, there are genes encoding: a GH2 β-glycosidase (according to CAZy often characterized as a galactosidase or glucuronidase), one or two GH16 glycoside hydrolases (based on previously characterized enzymes from marine bacteria putatively harbor either endoglucanase or agarase activity), and a GH149 enzyme, which, according to the CAZy database, can act as a β-1,3-glucan- or oligoglucan phosphorylase (cazy.org, visited 19-01-2026). These upstream genes are conserved in most of the investigated clusters. However, in some clusters, the GH149 enzyme is replaced by a GH3 enzyme. These genes are all predicted to encode enzymes important for converting marine biomass polymers, although not necessarily only laminarin conversion, considering the gene content in the complete PUL. In many of the gene clusters, the first gene is annotated as LuxR, a transcriptional regulator, indicating that the genes indeed are coregulated, as in a PUL. Moreover, hypothetical genes or genes with unknown functions are identified that may be involved in protein-carbohydrate interactions, such as the PKD-domain that, based on its predicted domain structure, is believed to be involved in such interactions (InterPro, IPR000601).

According to predictions by the SignalP and LipoP servers and subcellular location prediction using CELLO, *Ml*GH17A was found to have a signal peptidase II cleavage site. This suggests that after cleavage by signal peptidase II, *Ml*GH17A would be anchored to the membrane through the sulfhydryl group of Cys, which is predicted to be Cys20. The enzyme is predicted to be in the periplasmic compartment of the cell. GH30_1 and *Ml*GH17B lacked detectable signal peptides and were predicted to be located in the cytoplasmic part of the cell. The fact that *Ml*GH17A is an enzyme involved in the apparent laminarin-specific gene cluster in the PUL from *M. lutaonensis* ISCAR-4703, with homologs at the corresponding position in all *Muricauda* species, makes it an interesting target for further characterization.

### Production and purification of *Ml*GH17A

*Ml*GH17A was produced in *E. coli* as an MBP-fusion protein and was purified using a two-step affinity chromatography process. In the first step, MBP-affinity chromatography was employed to purify the fusion protein MBP-Smt3-*Ml*GH17A-6×His-tag. However, during this step, spontaneous cleavage of the MBP protein was observed, as indicated in Lane 1 of supplementary Fig. S1. In order to cleave the remaining attached MBP, the recombinant protein was incubated with the Ulp1 protease at 30 °C for 1 h (Lane 2, supplementary Fig. S1) and subsequent affinity purification using a His-Trap column resulted in a more than 90% pure protein (Lane 3, supplementary Fig. S1) that exhibited storage stability at 4 °C and a low tendency to aggregate. The SDS-PAGE analysis confirmed the level of purity achieved for the enzyme.

The estimated molecular weight of the construct, including MBP-Smt3-*Ml*GH17A-6×His-tag, was calculated to be approximately 103 kDa. This comprises the molecular weight of *Ml*GH17A, which is 48 kDa, combined with the molecular weight of the MBP-Smt3 domain and the 6×His-tag, calculated to be 55 kDa.

### Activity optimum determination

The activity of the enzyme was first tested at room temperature at different pHs (ranging from pH 3 to 9) using Thin Layer Chromatography (TLC). After screening of the products, a narrower pH range of 3 to 6 was selected, and the activity was measured at temperatures ranging from 13 °C to 53 °C. The results indicated that the enzyme exhibited the most significant activity at a pH close to 4. By narrowing the pH range, it was observed that *Ml*GH17A exhibited its highest activity at pH 4.5 and at a temperature of 42 °C (Fig. 2). Based on this data, a temperature of 40 °C and pH 4.5 were selected as optimal conditions for further experiments.

**FIG 2.**
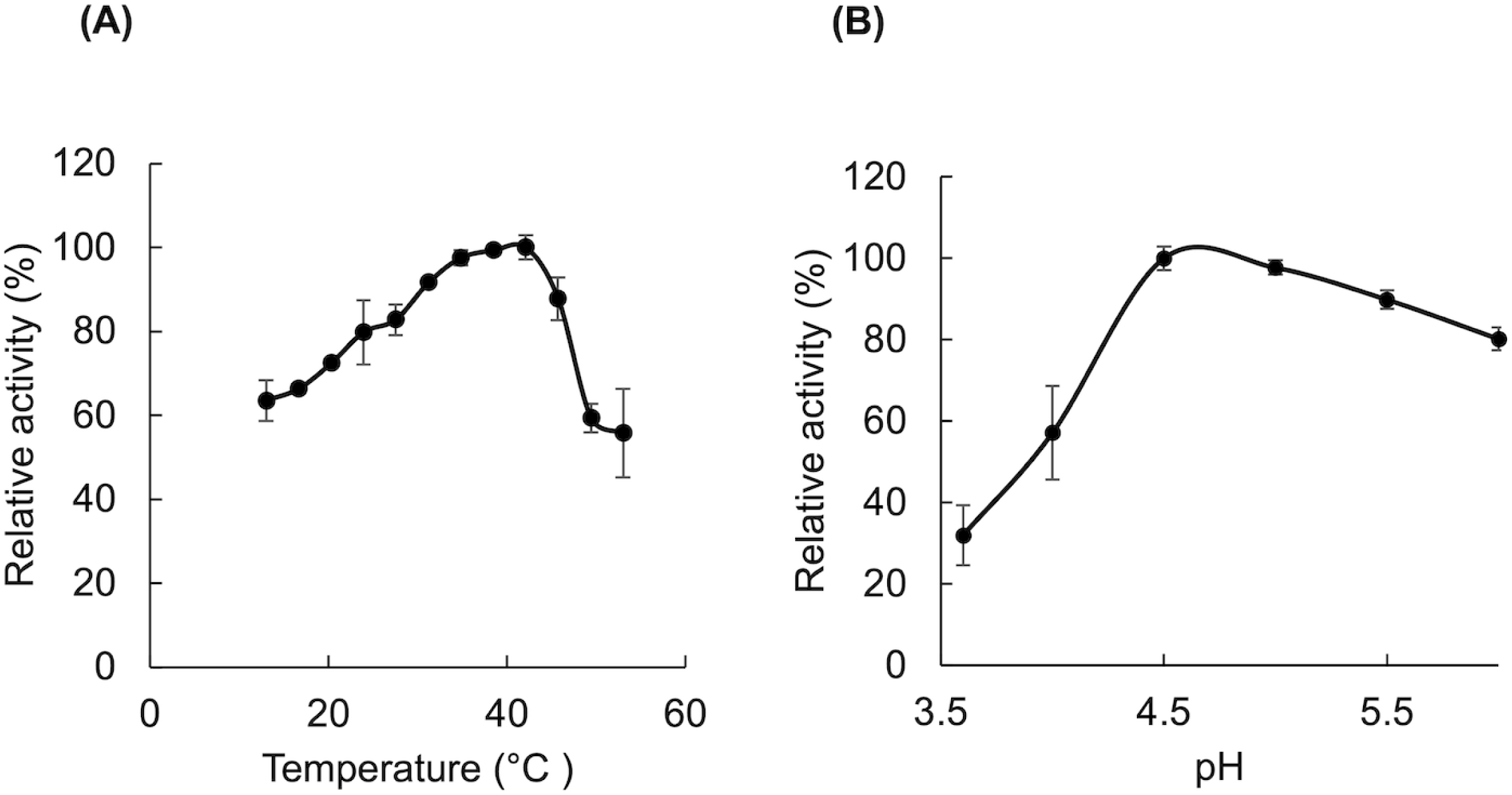
pH and temperature optima of *Ml*GH17A. The relative activity of the enzyme at pH 4.5 and various temperatures is shown in (A) and the relative activity of the enzyme at 42 °C in various pHs is represented in (B). All measurements were conducted in triplicates, and the results include the mean value and standard deviation.

### Melting temperature determination

NanoDSF thermogram of *Ml*GH17A (supplementary Fig. S2) shows a T_m_ of the enzyme, representing the inflection temperature of thermal unfolding, of 59 ± 0.1 °C, and a T_onset_ value, representing the onset temperature of thermal unfolding, of 52.4 ± 0.5 °C. From the observations, it is evident that the temperature optimum for activity (T_opt_ = 42 °C) was lower than the unfolding temperature (59 °C), and approximately 10 °C lower than the T_onset_ (52.4 °C). Differences between the temperature for the activity optimum and thermal unfolding were also found for the transglycosylase *Ml*GH17B (16), encoded in the same PUL. It was also noticed that an increase in temperature from 40 °C (T_opt_) to 53 °C (T_onset_) resulted in activity loss of around 45% (prior to reaching T_m_). This behavior might be attributed to subtle changes in the enzyme’s structure, such as local unfolding (without exposure of residues contributing to the detector signal) or changes in the flexibility of loops, which play a crucial role in the catalytic activity of enzymes with the TIM-barrel (β/α)_8_ structure. Such changes may reduce the enzyme’s ability to bind substrates or catalyze the reaction efficiently, resulting in activity loss prior to unfolding. The bumpy baseline in the first derivative graph (Fig. 3) supports and confirms this phenomenon, indicating that structural changes or disruptions occur in the enzyme.

**FIG 3.**
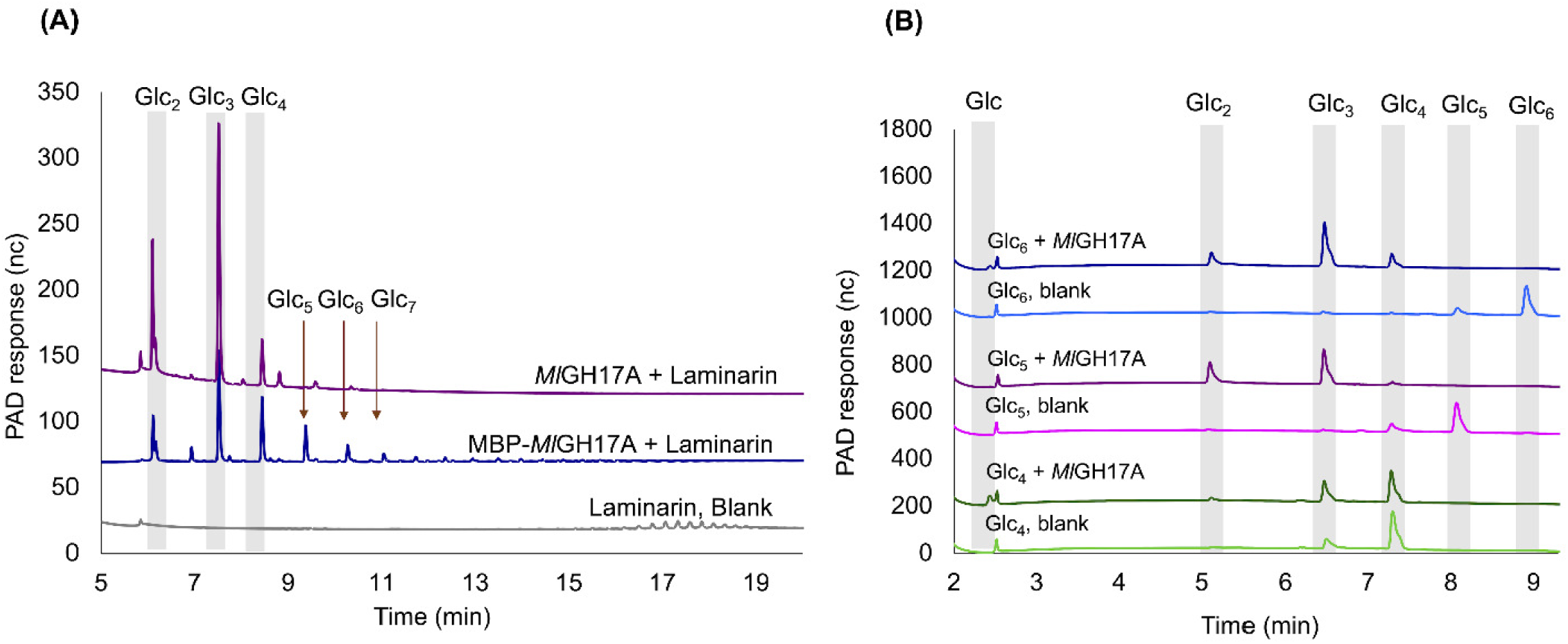
Product profile of *Ml*GH17A. Reaction products were obtained after 24 h incubation with (A) laminarin and (B) laminari-oligosaccharides, including laminaritetraose (Glc_4_), laminaripentaose (Glc_5_), and laminarihexaose (Glc_6_), at 40 °C and pH 4.5. The elution time of laminari-oligosaccharides standards (Glc_2-7_) is shown by gray bars and arrows. MBP-*Ml*GH17A represents MBP fused *Ml*GH17A.

### Substrate specificity and product profiles

Screening of hydrolysis products was made by HPAEC–PAD. No significant activity was found on the substrates lichenan, amylose, pustulan, or pullulan (data not shown), while the enzyme displayed activity on laminari-oligosaccharides and laminarin (Fig. 3), showing that the enzyme preferentially cleaves polysaccharides with β-1,3 linkages, including laminarin and laminari-oligosaccharides. Incubation of *Ml*GH17A with laminarin resulted in release of a range of oligomers, with the main products being Glc_2_, Glc_3,_ and Glc_4_ (DP2-4) (Fig. 3A). No activity was observed on laminaribiose (Glc_2_) or laminaritriose (Glc_3_), while minimal activity was detected on laminaritetraose (Glc_4_) (Fig. 3B). Activity on Glc_4_ was, however, considerably lower than the activity observed on laminaripentaose (Glc_5_) and laminarihexaose (Glc_6_) that were completely converted under the conditions used (Fig. 3B). The activity of the MBP-fusion protein (MBP-*Ml*GH17A) on laminarin substrates was also evaluated (Fig. 3A) to assess the potential for eliminating the second-step purification in industrial applications. The analysis revealed that the activity was notably lower in comparison to *Ml*GH17A without the MBP protein, and products with a wide range of degrees of polymerization (DPs) were produced.

To confirm endo-activity, the enzymatic reaction was performed with a very low concentration of the enzyme using laminarin as the substrate, and with sampling after short intervals (10-30 seconds). The results showed that the intensity of long-chain oligosaccharides progressively decreased over time, indicating conversion to shorter-chain oligosaccharides, specifically formation of Glc_2_, Glc_3_, and Glc_4_ (Supplementary Fig S3). The HPAEC-PAD results clearly show that Glc_2_ is not the only product. Glc_3_ and Glc_4_ were also produced, further demonstrating the endo-activity of the enzyme, as it cleaved internal glycosidic bonds within the oligosaccharides, rather than acting at the chain ends.

The distribution of products in the reaction mixture, utilizing laminari-oligosaccharides and laminarin as substrates, is shown in Table 1. As previously mentioned, Glc_4_ is not a favorable substrate for *Ml*GH17A, with only 54 % of the substrate being hydrolysed during the course of the reaction, whereas Glc_5_ – Glc_8_, and laminarin were completely hydrolysed. Glc_3_ was a predominant product, reaching close to 50% of the reaction products, with the remaining portion being evenly distributed between Glc_2_ and Glc_4_ (Table 1), highlighting the enzyme’s consistent behaviour toward substrates with a degree of polymerization above DP6.

**TABLE 1.** Relative abundance of the products of *Ml*GH17A in the reaction mixture utilizing laminari-oligosaccharides and laminarin as substrates.

| Substrate | Glucose | Glc<sub>2</sub> | Glc<sub>3</sub> | Glc<sub>4</sub> |
| --- | --- | --- | --- | --- |
| Laminaritetraose (Glc<sub>4</sub>) | 21.8 ± 3.73 | 4.1 ± 0.72 | 28.32 ± 0.89 | 45.79 ± 4.23 |
| Laminaripentaose (Glc<sub>5</sub>) | 7.78 ± 0.53 | 35.91 ± 0.23 | 50.25 ± 1.05 | 5.98 ± 0.5 |
| Laminarihexaose (Glc<sub>6</sub>) | 6.11 ± 1.06 | 17.45 ± 2.61 | 54.72 ± 1.16 | 21.72 ± 0.44 |
| Laminariheptaose (Glc<sub>7</sub>) | 7.7 ± 0.93 | 19.94 ± 0.42 | 51.14 ± 0.55 | 21.22 ± 0.1 |
| Laminarioctaose (Glc<sub>8</sub>) | 6.43 ± 0.31 | 25.38 ± 0.64 | 48.51 ± 0.84 | 19.68 ± 0.21 |
| Laminarin | 8.27 ± 0.17 | 25.98 ± 0.26 | 46.64 ± 0.1 | 19.11 ± 0.28 |

The synergy effect of *Ml*GH17A and the previously characterized *Ml*GH17B, located in the same cluster and isolated from the same strain (10), was also studied (Fig. 4). *Ml*GH17B is a β-1,3-glucanosyltransglycosylase that cleaves off two residues from the reducing end of the substrate and transfers the remaining part to another substrate molecule via a β-1,6 linkage, thereby synthesizing branched (if the linkage is to an internal second or third glucose from the non-reducing end of the acceptor) or kinked (if the linkage is to the non-reducing end glucose of the acceptor) oligosaccharides. In this study, it was noted that the branched oligosaccharides generated by *Ml*GH17B could be utilized by *Ml*GH17A, indicating that the enzyme can accommodate β-1,6 branched substrates in its active site. The products generated by *Ml*GH17A (DP2-4) could, however, not serve as substrates for *Ml*GH17B due to the fact that *Ml*GH17B exhibits very low activity towards substrates with a DP lower than DP5.

**FIG 4.**
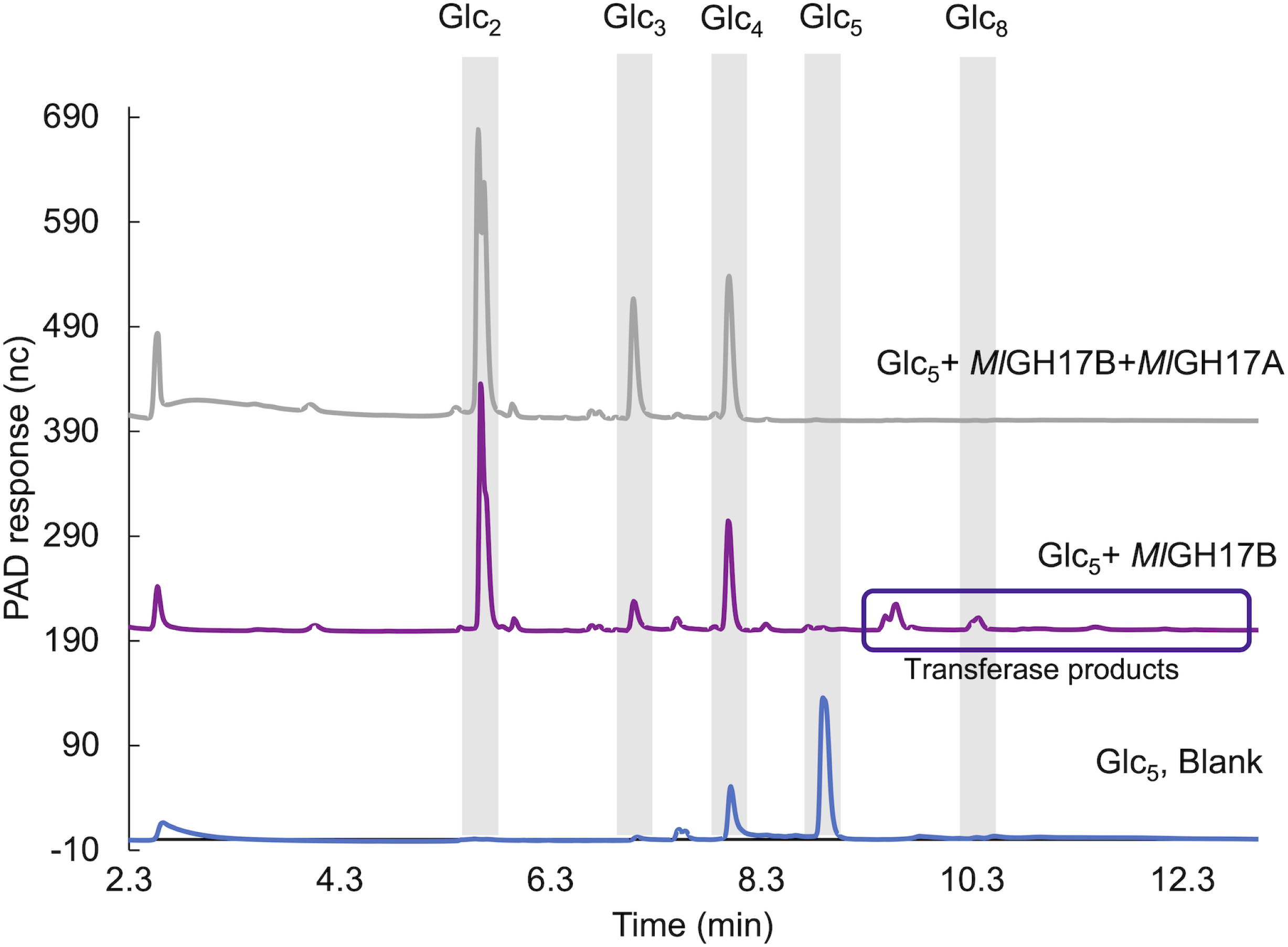
Sequential incubation of Glc_5_ with *Ml*GH17B (β-1,3-glucanosyltransglycosylase) and then with *Ml*GH17A. The reactions were performed at pH 6.0 (pH optima of *Ml*GH17B) with an intermediate boiling.

This result was confirmed by NMR and MALDI-TOF analysis of the products from the enzymatic reactions. NMR of Glc_8_ (produced by *Ml*GH17B) showed β-1,6 linkages on C2/C3 from the non-reducing end, with no kinked structures detected. After incubation with *Ml*GH17A, MALDI-TOF and NMR confirmed the formation of Glc_3_ and Glc_5_, demonstrating cleavage of β-1,6 linkages (Fig. 5).

**FIG 5.**
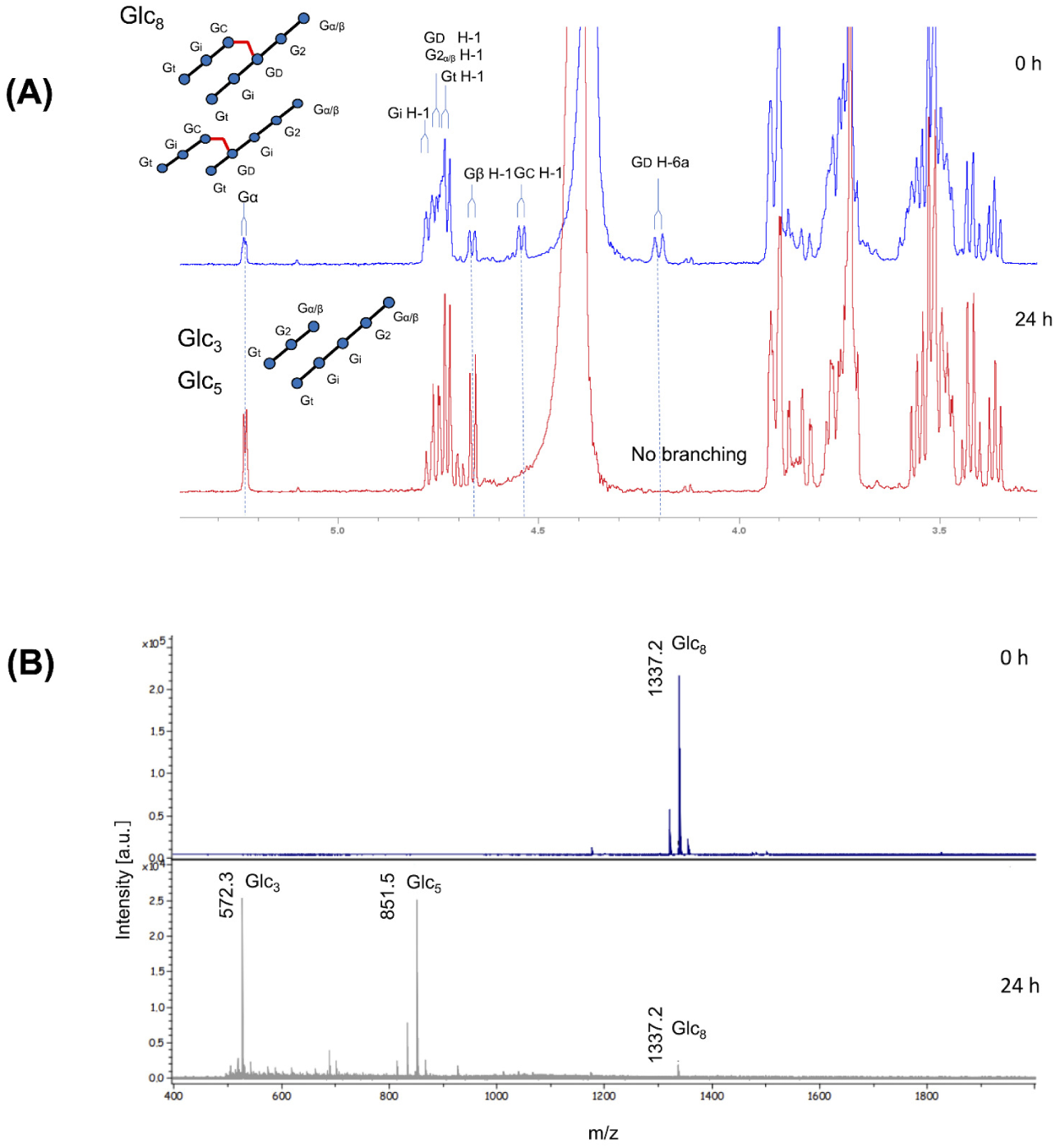
NMR and MALDI_TOF analysis of the products from synergy reaction of *Ml*GH17A and *Ml*GH17B. (A) 1D ^1^H NMR spectra of branched branched-octasaccharide before and after incubation with *Ml*GH17A at 311K in NMR tube. (B) MALDI-TOF MS spectra of branched octasaccharide before (0 h) and after (24 h) incubation with *Ml*GH17A at 311K in NMR tube. The possible linkages are shown as Gt [β-D-Glc*p*-(1→3)-]; Gi [-(1→3)-β-D-Glc*p*-(1→3)-]; G2 (next to reducing end) [-(1→3)-β-D-Glc*p*-(1→3)-]; Gα/β (reducing end) [-(1→3)-β-D-Glc*p*]; GC [-(1→3)-β-D-Glc*p*-(1→6)-]; and GD [-(1→3,6)-β-D-Glc*p*-(1→3)-].

### Determining kinetic parameters

In order to compare the catalytic efficiency of *Ml*GH17A with previously characterized laminarinases, the kinetic parameters were determined by quantification of the reducing ends produced after incubation of the enzyme with laminarin (Supplementary Fig. S4). The kinetic parameters of the enzyme were found to be *K_m_* 0.65 mM, *k*_cat_ 5.77 s^-1^, and *k*_cat_/*K_m_* 8843 M^-1^s^-1^. Kinetic comparisons were made between *Ml*GH17A and two characterized GH17 laminarinases: *Fb*GH17A, a laminarinase from a *Formosa* sp. Hel1_133_33 strain, (1) and *Vb*GH17A, a laminarinase from *Vibrio breoganii* (17). *Fb*GH17A exhibited a similar product profile as *Ml*GH17A, producing a mixture of laminari-oligosaccharides and glucose, although glucose was a minor product for *Ml*GH17A. In contrast, *Vb*GH17A produced oligosaccharides with higher DP (DP>3). It is noteworthy that the substrate used for kinetic determination of all three enzymes was laminarin from *L. digitata* (Sigma-Aldrich (Merck)).

The kinetic parameters were evaluated for *Fb*GH17A at pH 7 and 37 °C, and for *Vb*GH17A at pH 6.5 and 30 °C (enzymes’ optimum conditions). The kinetic parameters of *Ml*GH17A were measured at pH 4.5 and 40 °C (optimum conditions). Considering the mentioned conditions, the *K_m_* value of *Ml*GH17A (0.65 mM) falls within approximately a similar range (although 2.5 times lower) as *Fb*GH17A (1.6 mM), but it is significantly higher than the *K_m_*value of *Vb*GH17A (0.52 µM). *Fb*GH17A displayed the highest turnover number (57 s^-1^), almost 10 times higher than the *k*_cat_ of *Ml*GH17A (5.7 s^-1^) and *Vb*GH17A (6 s^-1^).

### Bioinformatic analysis

#### Laminarinases in the GH17 family

Characterized laminarinases can be found in five glycoside hydrolase families (16, 17, 55, 64, and 81) according to the CAZy database (cazy.org, visited 19-01-2026) (15, 18) and each family has at least one structure determined member. Three of the families GH55, GH64, and GH81 include inverting laminarinases with structures displaying a parallel β-helix structure, a crescent-like fold structure, and an (α/α)_6_-barrel structure, respectively. Two GH families, GH16 and GH17, include retaining laminarinases. GH16 contains β-jelly roll folded laminarinases, where two glutamates function as a proton donor and a nucleophile. Enzymes belonging to GH17 share a (β/α)_8_-barrel fold with two glutamates acting as the proton donor and nucleophile and include four bacterial characterized endo-acting and one exo-acting laminarinases. *Fa*GH17A is an endo-acting glycoside hydrolase from *Formosa agariphila* (GenBank CDF79584.1) with high specificity towards β-1,3-linked glucans (18). Two enzymes were isolated from *Formosa* sp. strain Hel1_133_33 (1); one debranched laminarin-specific exo-acting β-1,3-glucosidase (*Fb*GH17B) (GenBank AOR29491.1), and the other (*Fb*GH17A) is an endo-acting β-1,3-glucan hydrolase (GenBank AOR29489.1). *Vv*GH17 is an endo-acting β-1,3-glucanase isolated from *Vibrio vulnificus* (GenBank ASM98089.1) producing disaccharides as the major product (19). Finally, the laminarinase *Vb*GH17A, isolated from *Vibrio breoganii* (GenBank OEF87991.1), is an endo-acting enzyme, producing oligosaccharides with the smallest DP of 4 from a laminarin substrate (17). A multiple sequence alignment of catalytic modules of *Ml*GH17A, the previously characterized GH17 laminarinases, and *Ml*GH17B (the β-1,3-glucanosyltransglycosylase from *M. lutaonensis* ISCAR-4703) is shown in Fig. 6 and the sequence identity matrix is shown in Table S1. The catalytic domain of *Ml*GH17A exhibited a relatively low sequence identity with the catalytic domain of the exo-acting *Fb*GH17B (21%, with a query coverage of 56%) and with the catalytic domain of *Ml*GH17B (24.4%, with a query coverage of 47%), while these two latter enzymes were similar to each other (Table S1) indicative of a different specificity group. This can also motivate presence of both enzyme types in the PUL (Fig. 1)

**FIG 6.**
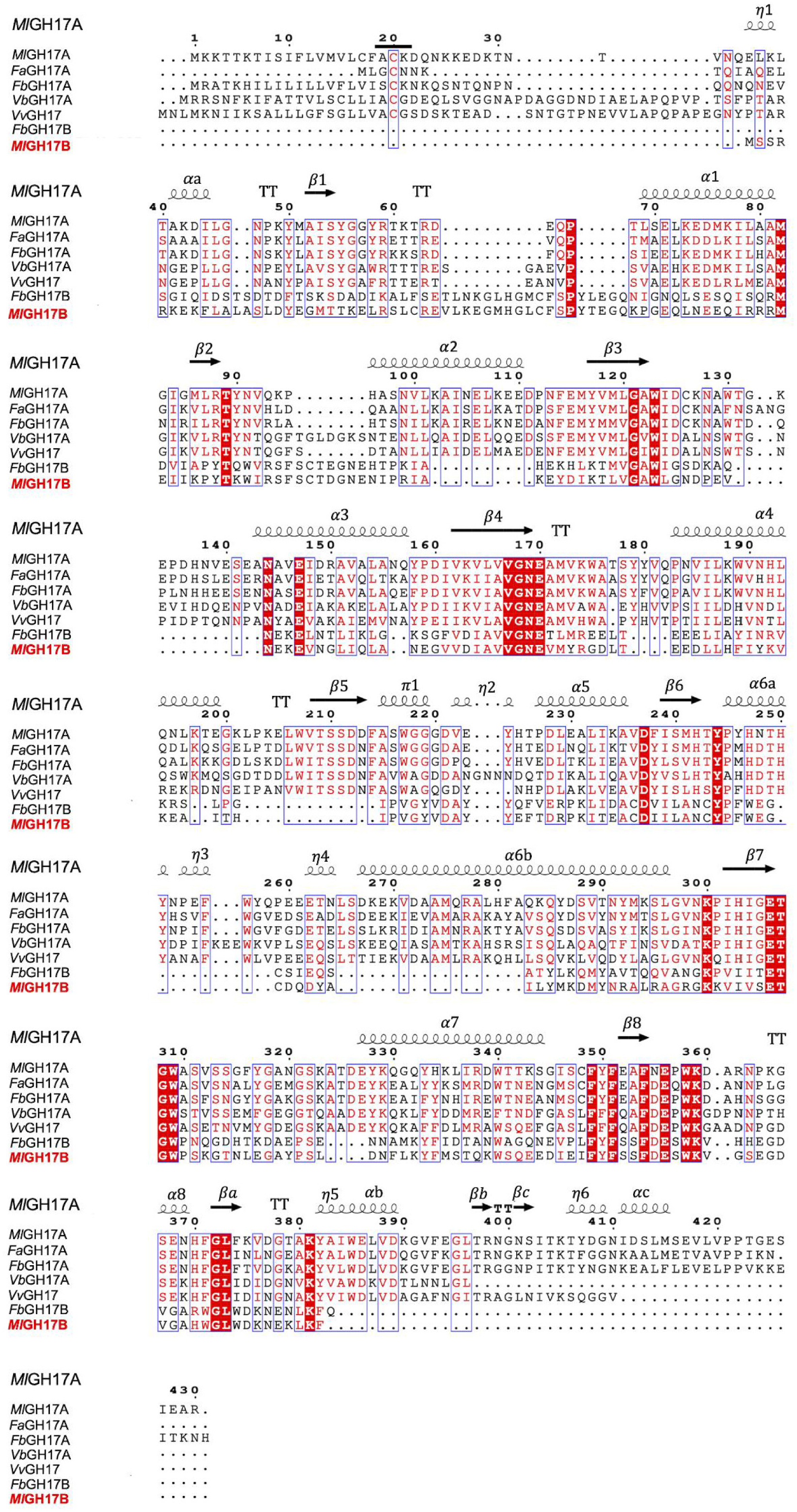
Multiple sequence alignment of characterized laminarinases from the GH17 family, laminarinase (*Ml*GH17A) and β-1,3-glucanosyltransglycosylase (*Ml*GH17B) from *M. lutaonensis* ISCAR-4703. *Fa*GH17A (GenBank CDF79584.1), laminarinase from *Formosa agariphila*; *Fb*GH17A (GenBank AOR29489.1), laminarinase from *Formosa* sp. Hel1_33_131; *Fb*GH17B (GenBank AOR29491.1), laminarinase from *Formosa* sp. Hel1_33_131; *Vb*GH17A (GenBank OEF87991.1), laminarinase from *Vibrio breoganii*; *Vv*GH17 (GenBank ASM98089.1), β-1,3-glucanase from *Vibrio vulnificus; Ml*GH17B (GenBank WIW39500), β- β-1,3-glucanosyltransglycosylase from *M. lutaonensis* ISCAR-4703. The predicted structure of *Ml*GH17A is illustrated on top of the alignment, indicating β sheets,α helices, 310-helix marked as η, strict β-turns marked as TT, and π-helices marked as π. For *Ml*GH17A, the predicted lipid anchoring cysteine (Cys20) is underlined and it is predicted that the signal peptidase II cleaves immediately before this cysteine.

The sequence identity with *Vb*GH17A was 45.4% (query coverage of 80%), while with *Vv*GH17, it was 49.7% (query coverage of 95%) and the identity to both *Fa*GH17A (66.5%, with a query coverage of 89%) and *Fb*GH17A (63.4%, with a query coverage of 90%) was even higher, showing that *Ml*GH17A shared significant sequence conservation with the endoglucanases.

In barley GH17 endoglucanases, an additional subdomain, extending the sequence and aglycone subsites, is present. This subdomain is reported to be missing in both bacterial and fungal transglycosylases (Supplementary Fig. S5), shortening their sequences and reducing the number of aglycone subsites (20, 21). It is, however, not directly related to the activity, as sequence analysis and structure modeling have shown this subdomain to be absent also in bacterial GH17 endo-β-glucanases (10) (Supplementary Fig. S5 and Supplementary Fig. S6), despite their slightly longer sequences compared to the transglycosylases (see Fig. 7). Notably, both the endo-glucanase *Ml*GH17A and the transglycosylase *Ml*GH17B in the PUL are of a shorter sequence type than the barley enzymes, indicative of fewer aglycone subsites.

**FIG 7.**
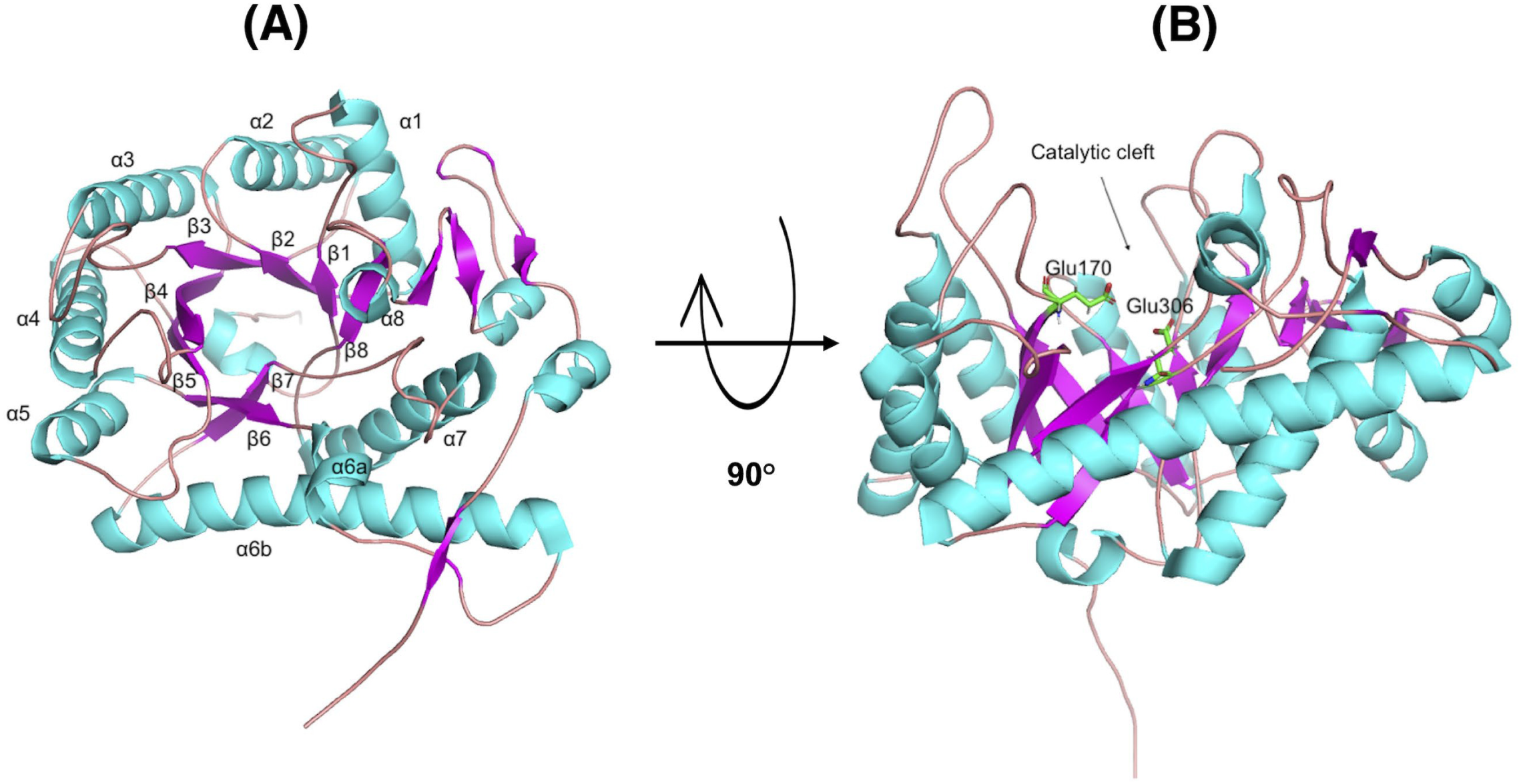
Homology model of *Ml*GH17A represented as a ribbon diagram. The TIM-barrel (β/α)_8_ structure of the enzyme is illustrated (A) from the front view and (B) from the side view. The catalytic residues (E170 and E306) are shown in green.

Compared to the *Formosa* enzymes, *Ml*GH17A shares its position in the PUL with *Fb*GH17A, and these enzymes are both endo-glucanases. *Ml*GH17B, on the other hand, is a β-1,3-glucanosyltransglycosylase releasing laminaribiose (10), while the gene at the corresponding position in the PUL in *Formosa* is encoding the exo-glucosidase *Fb*GH17B. The high sequence similarity between *Fb*GH17B and *Ml*GH17B (53% seq. id, Table S1), however makes it likely that they have related functions in their respective microorganism, despite the difference in activity profiles.

#### Three-dimensional modeling of MlGH17A

So far, among the bacterial GH17 laminarinases, the only successfully determined crystal structure is of *Fb*GH17A, a laminarinase from *Formosa* sp. Hel1_133_33 (1). To generate a homology model of *Ml*GH17A, the crystal structure of *Fb*GH17A (63% sequence identity and a query coverage of 90% with *Ml*GH17A) was used as the main template for modelling. The model validation gave an overall Z-score of −0.278 by YASARA software, as the weighted averages of the individual Z-scores using the formula Overall = 0.145 Dihedrals + 0.390 Packing1D + 0.465 Packing3D. According to VERIFY3D, which assessing the quality and compatibility of protein structure models with their corresponding amino acid sequences, 84.08% of the residues have averaged 3D-1D score>=0, indicating a good quality model (value about 80% indicates good quality model). ERRAT calculated the overall quality factor of the model as 98.98%. This score represents the quality of the model on a scale from 0 to 100, with higher scores indicating better agreement between the model and known protein structures. A good high-resolution structure typically produces values around 95% and higher. Ramachandran plot obtained by PROCHECK illustrated 93% of the residues are in the most favoured region, and 7% are located in the allowed region.

The three-dimensional model of *Ml*GH17A exhibited a TIM-barrel (β/α)_8_ structure, which is the common fold of all GH17 enzymes, and within this structure Glu170 functions as the proton donor, while Glu306 acts as the nucleophile (Fig. 7). As can be seen in Fig. 7 and Fig. 8, Glu170 and Glu306 are located at the C-terminal of β4 and β7, respectively.

**FIG 8.**
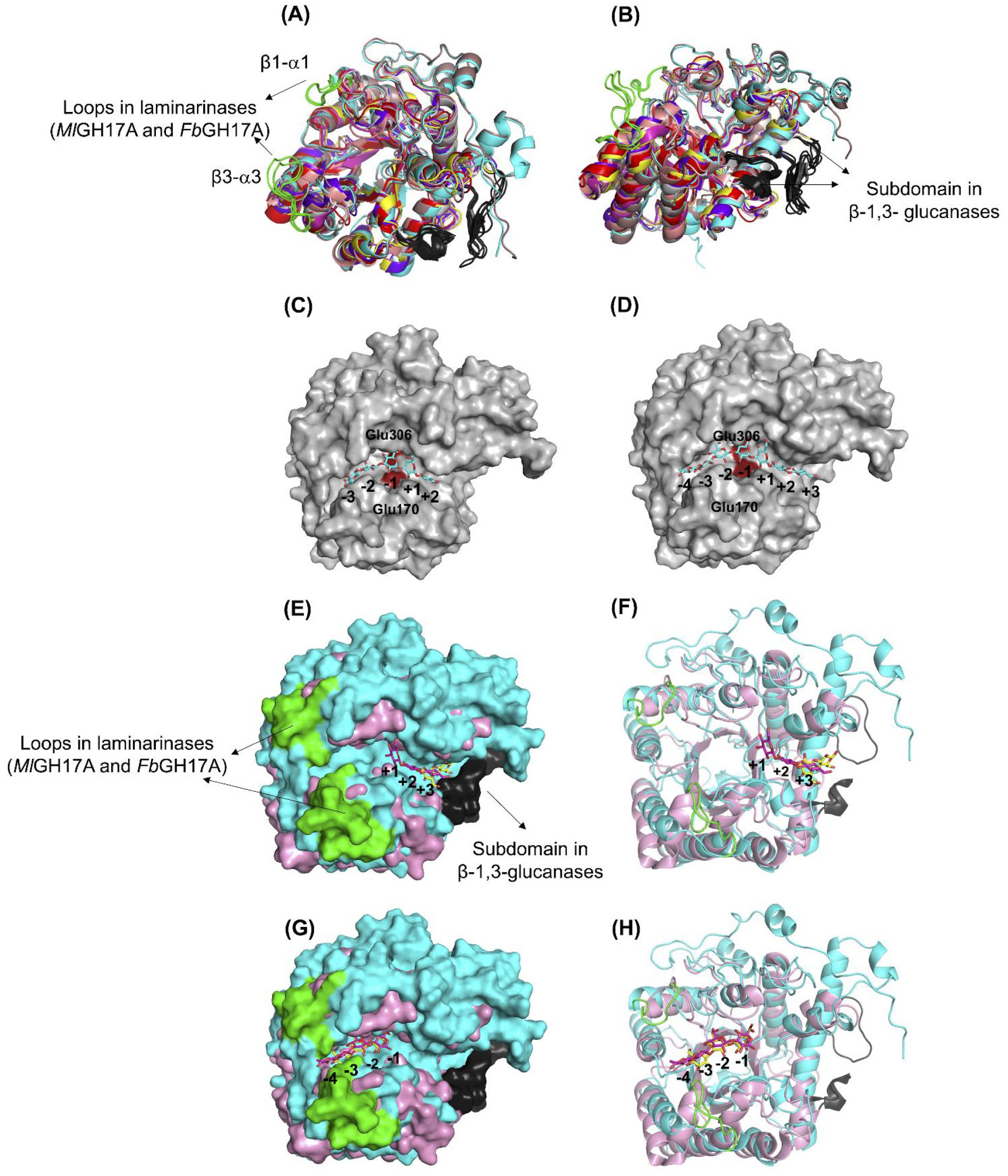
(A, B) Comparison of the three-dimensional model of *Ml*GH17A (cyan) with the crystal structure of GH17 plant β-1,3-glucanases such as endo-β-1,3-glucanase from *Musa acuminata* (PDB 2CYG, red), endo-β-1,3-glucanase GII (Bgl32) from *Hordeum vulgare* (PDB 1GHS, magenta), barley β-1,3-1,4-glucanase from *Hordeum vulgare* (PDB 1AQ0, yellow), endo-β-1,3-glucanase from *Solanum tuberosum* (PDB 3UR7, salmon), β-1,3-glucanase from *Hevea brasiliensis* (PDB 4HPG, gray), and β-1,3-glucanase from *Cryptomeria japonica* (PDB 6JMS, purple), and *Fb*GH17A, laminarinase from *Formosa* sp. Hel1_33_131 (PDB 6FCG, brown). The subdomain available in β-1,3-glucanases is shown in black, while the β1-α1 and β3-α3 loops of *Ml*GH17A and *Fb*GH17A are shown in green. (C, D) *Ml*GH17A docked to Glc_5_ and Glc_7_ ligand; the ligand is shown in blue and the catalytic residues E170 and E306 are shown in red. In sillico model of MlGH17A shows a narrow, extended substrate binding cleft to bind β 1,3 glucan chain of at least three monomers without space for β 1,6 side chains. (E, F) and (G, H) comparison of the aglycone and glycone subsites of *Ml*GH17A (cyan) and GluB20-2 (PDB 4GZJ, pink); the subdomain of Glu20-2 are shown in black while the β1-α1 and β3-α3 in *Ml*GH17A are shown in green; the ligand of Glu20-2 is shown in yellow and for *Ml*GH17A is shown in magenta.

Superimposition of *Ml*GH17A with structure determined and modelled candidates in GH17 showed some interesting differences. In bacterial laminarinases (or β-1,3-glucanases) such as *Ml*GH17A, *Fb*GH17A (PDB 6FCG), *Fa*GH17A, *Vb*GH17A and *Vv*GH17, the β3-α3 loop is significantly longer compared to that in the bacterial transglycosylases, fungal *Rm*Bgt17A, and debranched laminarin specific exo-acting β-1,3-glucosidase *Fb*GH17B (supplementary Fig. S5 and Fig. S6) and *Ml*GH17B. This suggests presence of additional glycone subsites in these enzymes, while the aglycone part of these enzymes is shorter than in the plant β-1,3-glucanases. Moreover, in all bacterial GH17 enzymes (including *Ml*GH17A) as well as the fungal *Rm*Bgt17A, the β1-α1 loop is longer than the corresponding loop in plant β-1,3-glucanases, however, the size and the orientation of this loop around the active site varies among the enzymes.

#### Docking of laminari-oligosaccharides

Substrate binding in the modeled structure of *Ml*GH17A was compared with that in the crystal structure of laminarinase *Fb*GH17A from *Formosa* sp. Hel1_33_131 (PDB 6FCG) and in the crystal structure of eukaryotic β-1,3-glucanases (Fig. 8A-8B). Docking experiments for *Ml*GH17A revealed that the active site of *Ml*GH17A comprises seven putative subsites: four glycone subsites (−4, −3, −2, and −1) and three aglycone subsites (+1, +2, and +3) (Fig. 8C-8D). The enzyme had a longer catalytic cleft in the glycone part compared to the plant β-1,3-glucanases (Fig. 8E-8H).

From the structural analysis of the β-1,3-glucanase, GluB20-2, from potato (*Solanum tuberosum*) and its mode of action, it is evident that substrate binding to glycone subsites (−3, −2 and −1) is crucial for the enzyme’s significant activity since several hydrogen-bonding interactions were detected between the amino acids and glucose moieties in the −1, −2 and −3 subsites. However, GluB20-2 lacks the crystal structure of the ligand-protein complex with DP4 ligand in the aglycone subsites due to the weak interactions of the glucosyl residues at the +1 and +2 subsites. Only laminaribiose is visible in the +3 and +4 subsites within the crystal structure. These weak interactions may allow the leaving group to depart from the catalytic cleft, which can then be replaced by a water molecule crucial for hydrolyzing the glycosyl-enzyme intermediate (22).

Laminaripentaose (Glc_5_), was docked manually to the active site of *Ml*GH17A from −3 to +2 subsites. Fig. 9 illustrates the potential hydrogen bonding interactions occurring between the hydrolyzed products: laminaribiose and laminaritriose, from a DP5 ligand, and the active site residues. Fig. 9A indicates that the glucose in the +1 subsite has the potential for multiple hydrogen bonding. Specifically, hydrogen bonds were observed between catalytic residue Glu170 and OH2 and OH3 of the glucose ligand and between Tyr246 and OH4 of the glucose in the +1 subsite.

**FIG 9.**
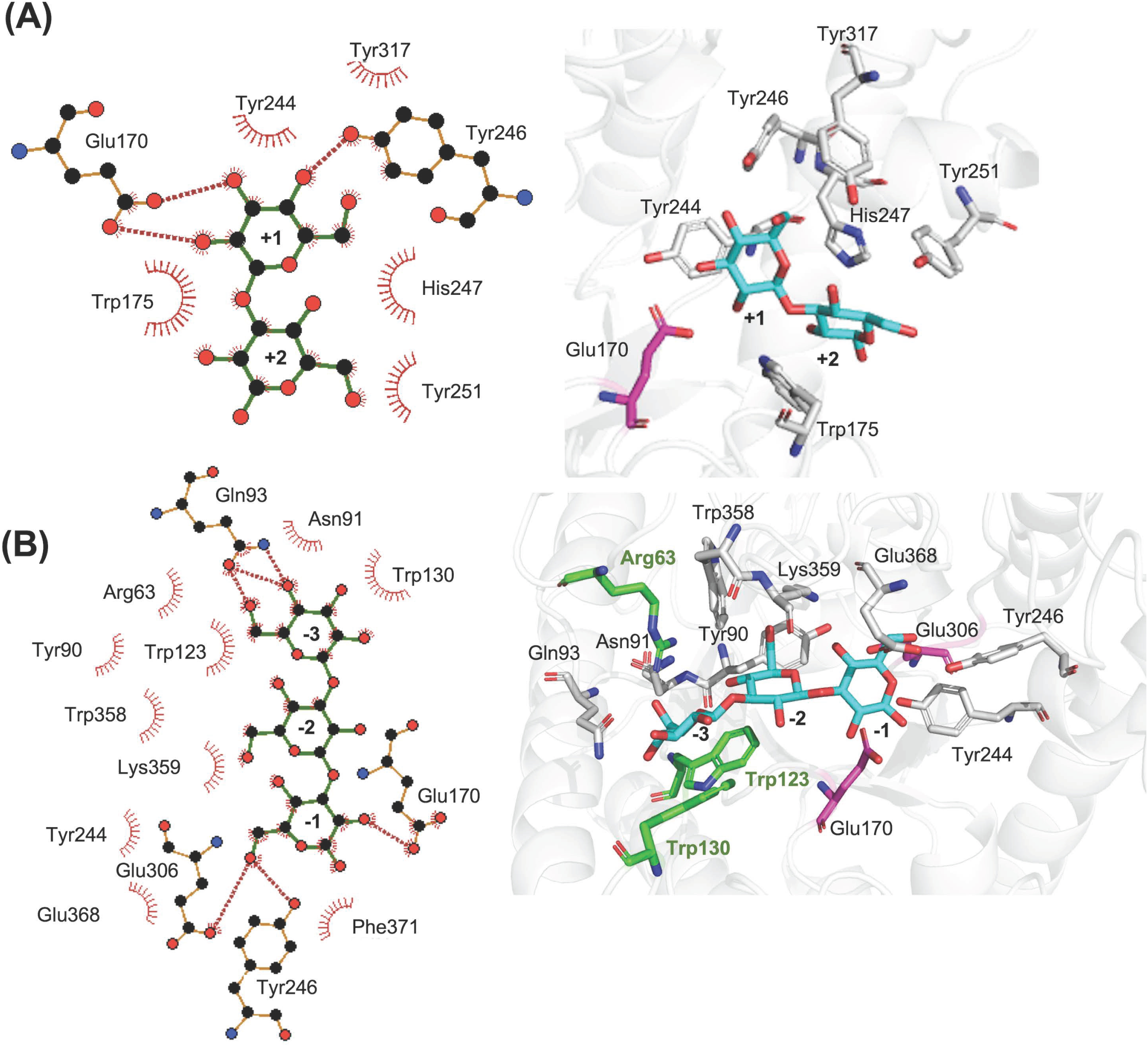
Laminaribiose and laminaritriose, products of enzymatic hydrolysis of Glc_5_, docked into (A) putative aglycone (+1 to +2) subsites and (B) putative glycone (−1 to −3) subsites of *Ml*GH17A. To the left, a 2D plot, obtained from Ligplot^+^, illustrates hydrogen bonds (Red dotted lines) and hydrophobic interactions (red stripes). To the right, 3D representation of the catalytic site (graphically presented via PyMOL) is shown, and the catalytic residues within 3Å are illustrated in gray while E170 and E306 are shown in magenta. The catalytic residues located in the loops β1-α1 (Arg63) and β3-α3 (Trp123 and Trp130) are shown in green.

Multiple potential hydrogen bonds were also detected in the glycone subsites (−1 to −3). Specifically, OH6 of the glucose in the −1 subsite could potentially interact with both Tyr246 and the nucleophile Glu306. Furthermore, the proton donor Glu170 could form a hydrogen bond with OH2 of the glucose moiety in the −1 subsite. Additionally, there were possible hydrogen bond interactions between Gln93 and OH4, and OH6 of the glucose moiety in the −3 subsite (Fig. 9B). It is important to note that hydrophobic stacking, observed for both glycone and aglycone subsites, together with the direct hydrogen bonds affects the binding affinity and substrate specificity of the enzyme.

By manually expanding the ligand, adding glucose moieties in both aglycone and glycone subsites, in a correct conformation, it was revealed that the enzyme may have seven putative subsites, three in the aglycone part (+1 to +3) and four glycone subsites (−1 to −4). Fig. 10 demonstrates the catalytic residues and the potential hydrogen bonds between the hydrolyzed products of Glc_7_ ligand and the enzyme. When the length of the ligand was expanded, a greater number of hydrogen bonds were observed in the aglycone subsites (+1 to +3) compared to the detected hydrogen bonding interactions in subsites (+1, +2) using the shorter ligand (Fig. 9A). The previously found potential hydrogen bonding interactions between the proton donor (Glu170) and OH2 of the glucose in the +1 subsite and between Tyr246 and OH4 were identified. In addition, bonding between Tyr246 and OH6 was found in the +1 subsite and Tyr317 could also form a hydrogen bond with OH6 in the +1 subsite (Fig. 10A). There was also one potential hydrogen bond detected between Asn248 and OH2 in the +3 subsite.

**FIG 10.**
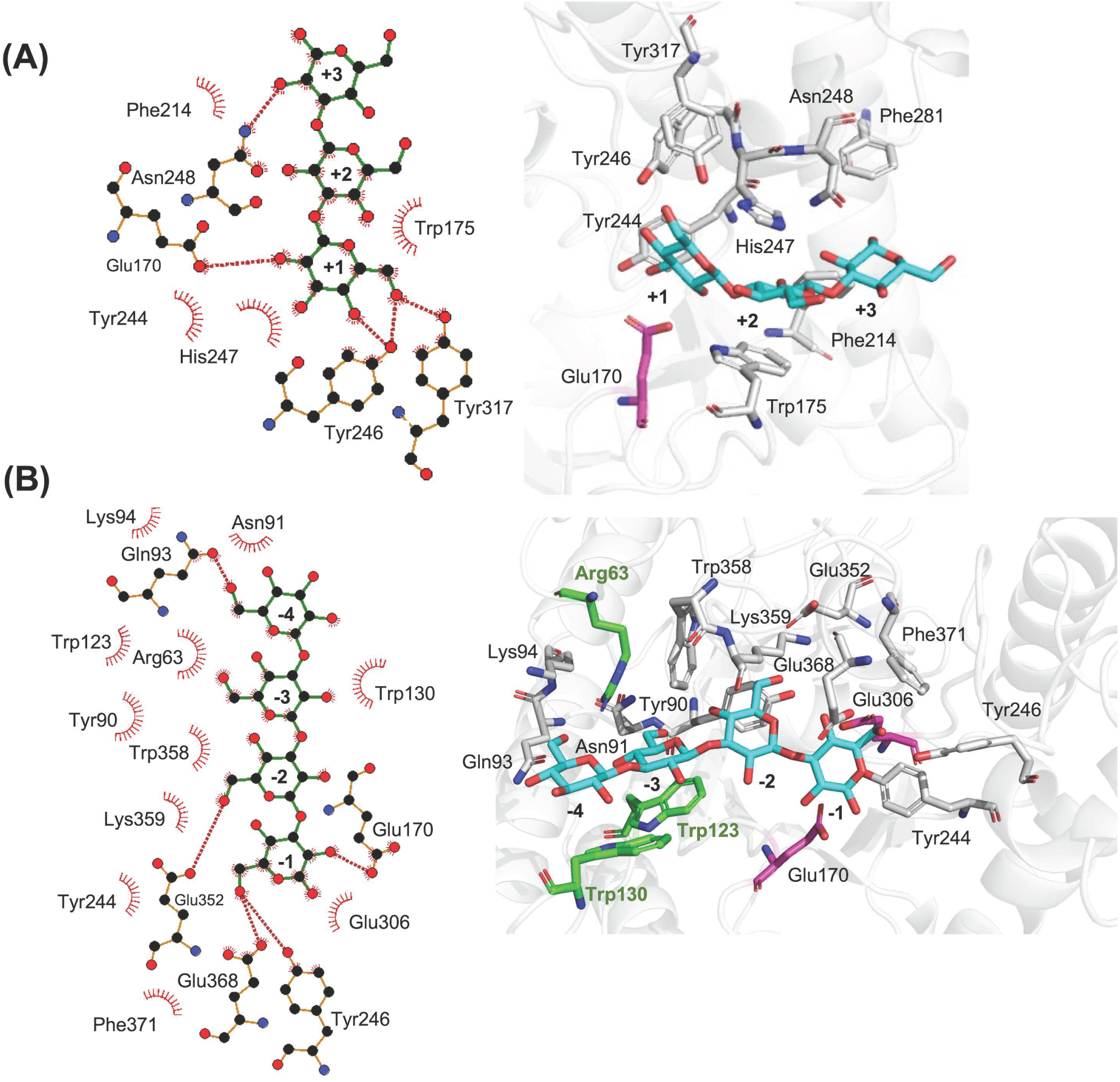
Laminaritriose and laminaritetraose, products of enzymatic hydrolysis of Glc_7_, docked into (A) putative aglycone (+1 to +3) subsites and (B) putative glycone (−1 to −4) subsites of *Ml*GH17A. To the left, 2D plot illustrates hydrogen bonds (Red dotted lines) and hydrophobic interactions (red stripes). To the right, a 3D representation of the catalytic site is shown, and the catalytic residues are shown in gray while E170 and E306 are shown in magenta. The catalytic residues located in loops β1-α1 (Arg63) and β3-α3 (Trp123 and Trp130) are shown in green.

In the glycone subsites, Tyr246 and Glu368 (instead of the potential nucleophile Glu 306) formed hydrogen bonds to OH6 in the −1 subsite and the proton donor Glu170 had a hydrogen-bonding interaction with OH2. In the −2 subsite, Glu352 hydrogen bonded to OH6, and in the −4 subsite Gln93 (which for the shorter ligand interacted in the −3 subsite) formed a hydrogen bond to OH6 of the glucose residue. (Fig. 10B). It can be seen that Arg63 which is located in the β1-α1 loop, Trp123 and Trp130 which are located in the β3-α3 loop are involved in hydrophobic interactions with glucose moieties in subsites −3 and −4 (Fig. 9B and Fig. 10B).

NMR and MALDI-TOF results indicate that the enzyme can cleave β-1,6 linkages. As shown in Fig. 8C and 8D, there is sufficient space at the −2 subsite to accommodate the linkage (for the product when the linkage is at C2 from the non-reducing end), as seen in Fig. 5A, considering ligand binding from the −3 to +2 subsites. However, bioinformatic analysis suggests that the enzyme may have a wider catalytic cleft with 7 subsites, implying that the ligand could bind at positions other than the −3 to +2 subsites, potentially accommodating β-1,6 linkages at C3 (Fig. 5A) from the non-reducing end in the −2 subsite.

## Discussion

The β-1,3 linked glucan polymer laminarin, is a storage carbohydrate in brown seaweed and other marine species that serves as a valuable source of energy and nutrients and is utilized by different species of marine bacteria. Marine bacterial species within the family *Flavobacteriaceae* have been shown to contain gene clusters encoding specific carbohydrate acting enzymes that can break down laminarin into glucose or oligosaccharides (1, 18), which can be used as an energy source or as building blocks for cellular processes. These gene clusters are organized in PULs, which are defined as strictly regulated, colocalized gene clusters encoding the enzyme and protein ensembles required for the saccharification of complex carbohydrates (7). Previous studies have identified PULs involved in the degradation of laminarin in different genera within the *Flavobacteriaceae* family such as *Formosa* (1, 18), *Gramella*, *Psychroserpens*, *Polaribacter*, as well as certain species within *Vibrio* family (23, 24). However, thus far, no information has been available regarding the laminarin utilization pathway in *Muricauda* species.

In the present study, PULs responsible for laminarin degradation were identified in various *Muricauda* species. These PULs exhibited a wide range of glycoside hydrolases and transglycosylases which highlights the ability of species within this genus to utilize strategies for laminarin degradation.

A conserved feature of all PULs detected in various *Muricauda* species is the presence of two genes encoding GH17 glycoside hydrolases, specifically an endo-acting laminarinase and a transglycosylase (β-1,3-glucanosyltransglycosylase in *Muricauda lutaonensis* ISCAR-4703), with genes encoding a GH30 (subfamily 1) glycoside hydrolase and an MFS transporter located in between. This characteristic was also observed in the PUL for laminarin utilization in *Polaribacter spp.* Hel1_33_49, Hel1_33_78, and Hel1_33_96 (25) as well as in *Formosa* sp. Hel1_33_131 (1). To date, no GH17 laminarinases or GH30 glycosyl hydrolases from the *Muricauda* genus have been characterized. Nonetheless, analysis of the PULs of other marine bacteria suggests that the GH30 (subfamily 1) enzyme is likely to be a β-1,6-glucosidase with the ability to degrade β-1,6 linkages. This characteristic was observed for *Fb*GH30 (GH30 β-glucosidase in the laminarin utilization pathway of *Formosa* sp. Hel1_33_13) which has 69% sequence identity (97% query coverage) to the one in the *M. lutaonensis* strain ISCAR-4703 PUL (1). The presence of a Major Facilitator Superfamily (MFS) transporter, located in the inner membrane, in the laminarin utilization pathway suggests a potential coupling between hydrolysis and product uptake processes (1). The enzyme, *Ml*GH17A, possesses a signal peptide, suggesting its likely location in the periplasm attached to the inner membrane of the bacteria. The enzyme exhibits the ability to act on both branched and unbranched substrates.

On the other hand, *Ml*GH17B lacks signal peptide, and is likely located in the cytoplasmic region of the cell. The genome of *M. lutaonensis* ISCAR-4703 encodes several intracellular glycosyltransferase family 2 (GT2) candidates (EC 2.4.1.34) which could be utilized by cells to synthesize β-1,3-glucan. The products of these enzymes can be used as substrates by *Ml*GH17B, which produces laminaribiose through hydrolysis. This laminaribiose can be utilized by cells for energy production. Additionally, long branched oligosaccharides which are produced by *Ml*GH17B through the transferase reaction can serve as storage carbohydrates. The presence of GH30 β-1,3-glucosidase and GH16_3 β-1,3-glucanase in the cytoplasm facilitates the utilization of *Ml*GH17B’s products by the cells.

The glycoside hydrolase enzyme from GH16 found at upstream locations in the same PUL is likely to be extracellular or membrane-bound. GH16 enzymes have been classified into a number of subfamilies in the CAZy database, and characterized enzymes from marine species include enzymes acting on many different glucans, not just laminarin (25), but also e.g., endo β-1,3(4)-glucanase as was observed in thermostable *Rhodothermus marinus* (25) or β-agarase activity as seen from a marine *Alteromonas* species (26).

The enzyme belonging to GH149 could be a β-1,3-glucan phosphorylase (cazy.org, visited 19-01-2026), aiding in degrading polysaccharides specifically β-1,3-glucan from external sources (27). This enzyme catalyzes the reversible degradation of β-1,3linked glucose polymers, utilizing inorganic phosphate from the environment. The presence of endo-acting GH16, GH2 β-1,6-glucosidase, and GH149 β-1,3-glucan phosphorylase, facilitate the breakdown of laminarin to oligosaccharides. These products can then be transported to the periplasmic part of the cell, broken down to oligosaccharides with a degree of polymerization not higher than 4, and then transferred to the cytoplasmic part of the cell for energy production or storage.

The endo-acting enzyme *Ml*GH17A, available in the laminarin utilization cluster of *Muricauda lutaonensis* strain ISCAR-4703, has demonstrated significant activity on laminari-oligosaccharides with DP5 and above, as well as on laminarin. The three-dimensional homology model of the enzyme reveals that the enzyme has a wide active site with seven putative subsites, −4 to −1 in the glycone part and +1 to +3 in the aglycone part, capable of accommodating β-1,6 branched carbohydrate molecules. This characteristic is particularly important because *Ml*GH17A is able to degrade laminarin which is a complex carbohydrate with occasional β-1,6 branches. The enzyme produces laminaribiose, laminaritriose, and laminaritetraose during hydrolysis, with laminaritriose being the dominant product. The homolog of *Ml*GH17A in *Formosa* sp. Hel1_33_13 (*Fb*GH17A), with 63% sequence identity, exhibits a corresponding endo-acting enzymatic activity, capable of acting on both branched and debranched laminarin, and is located in a corresponding position in a PUL of in the *Formosa* strain (1).

The homolog of *Ml*GH17B (*Fb*GH17B) is, however, not proposed to have a corresponding activity, despite its location at a similar position in the *Formosa* PUL. The enzyme is proposed to be an exo-acting β-1,3-glucosidase (1), hydrolyzing debranched laminarin, releasing glucose as the product. Unless a transglycosylase function of this enzyme is yet to be discovered, its catalytic mechanism thus differs from that of *Ml*GH17B. The products of *Ml*GH17B enzyme which are β-1,6 branched or kinked oligosaccharides can be stored as an energy reserve in the cell. It is noteworthy to mention that these two enzymes in the PUL (*Ml*GH17A and *Ml*GH17B) and their correspondence in the *Formosa* PUL (*Fb*GH17A and *Fb*GH17B) share only around 20% sequence identity, which confirms their distinct activities and cellular locations. It is worth noting that three residues (Tyr136, Trp191, and Glu192), previously identified as crucial for the transglycosylation activity of *Ml*GH17B (10), are not conserved in *Ml*GH17A and are replaced by Val173, His248, and Asn248, respectively (Fig. 6), which could be another indication of the distinct activity of these enzymes.

## Conclusion

This study revealed that the polysaccharide utilization locus (PUL) responsible for laminarin degradation in *Muricauda* species comprises a collection of glycoside hydrolase and transferase enzymes, enabling the bacteria to effectively uptake laminarin. Among these enzymes, a novel GH17 hydrolase, encoded by a gene located in the PUL in *M. lutaonensis* strain ISCAR-4703, was identified and characterized. This enzyme, designated *Ml*GH17A, is an endo-acting enzyme with the capability to degrade both branched and unbranched polysaccharides and oligosaccharides with a degree of polymerization (DP) of 5 and higher, producing laminaribiose (Glc_2_), laminaritriose (Glc_3_) and laminaritetraose (Glc_4_) as the main products. This enzyme is proposed to play a significant role in the degradation and utilization of laminarin by these bacteria. The bioinformatics analysis has identified that the enzyme possesses seven putative subsites: four glycone subsites (−4, −3, −2, and −1) and three aglycone subsites (+1, +2, and +3). Additionally, it was found that bacterial GH17 enzymes, such as laminarinases/β-1,3-glucanases and transglycosylases, lack the subdomain typically found in the aglycone part of GH17 plant β-1,3-glucanases. This absence results in a shorter active site in the aglycone region of the GH17 bacterial enzymes. However, it’s noteworthy that GH17 bacterial laminarinases/ β-1,3-glucanases may exhibit a longer active site in the glycone region (as observed for *Ml*GH17A); this extension could be attributed to the presence of the β3-α3 loop which is much shorter in GH17 eukaryotic enzymes and GH17 bacterial transglycosylases.

## Materials and methods

All materials were purchased from Sigma Aldrich unless otherwise specified. Laminari-oligosaccharides were supplied by Megazyme (Neogen).

### Identifying gene clusters in *Muricauda* species

Genome sequencing and annotation of the strain ISCAR-4703 was previously described by Allahgholi et al. (10). The amino acid sequences of *Ml*GH17A and a previously characterized β-1,3-glucanosyltransglycosylase, *Ml*GH17B, from the *Muricauda lutaonensis* ISCAR-4703 strain, were subjected to a BLAST search using the blastp tool in the NCBI BLAST suite (12), against the non-redundant protein sequence database. *Muricauda* species which contain the homologous of both enzymes (Table S2 and Table S3) with minimum query coverage of 90% and minimum sequence identity of 60% were selected for detection of the gene cluster involved in laminarin utilization by the bacteria. The reordered assembled draft genomes were obtained from GenBank and added to the RAST Server (Rapid Annotation using Subsystem Technology) (rast.nmpdr.org/) for annotation (28).

### Cloning, protein production, and purification

The sequence encoding the GH17 enzyme *Ml*GH17A was isolated from the genome of *M. lutaonensis* ISCAR-4703 strain (10), isolated from Yngingarlindir, an intertidal geothermal site located near the Reykjanes peninsula in Iceland.

A modified dual-tag pJOE system (29) was utilized to produce the recombinant protein with Maltose binding protein (MBP) domain at the N-terminus and His-tag at the C-terminus, separated by a *Saccharomyces cerevisiae* ubiquitin-like protein (Smt3) motif.

The plasmid of *Ml*GH17A was transformed into *E. coli* NiCo21 (DE3) strain and then cultured in shake flasks using LB-Lennox medium at 37 °C. Heterologous expression was induced when the optical density of the culture reached OD_600_ 0.5. The induction was done by adding L-rhamnose, to a final concentration of 0.25% (w/v), followed by 4 h incubation at 37 °C. The cells were subsequently harvested and lysed in binding buffer (20 mM Tris-HCl, pH 7.4, 200 mM NaCl) using a UP400S homogenizer. The resulting lysate was collected by centrifugation at 21,000×g and filtered through a 0.2 μm pore size filter.

The recombinant protein, MBP-Smt3-*Ml*GH17A-6×His, was purified by affinity chromatography using a 5 ml MBP-Trap HP column (GE Healthcare Life Sciences), utilizing the ÄKTA start FPLC purification system (GE Healthcare Life Sciences). Elution was performed using a maltose gradient of 10 mM in binding buffer over 10 column volumes. Fractions containing the recombinant protein were combined and stored at 4 °C for further purification.

The fractions containing the recombinant protein (MBP-Smt3-*Ml*GH17A-6×His) were combined and then incubated with Ulp1 (Ubiquitin-like-specific protease 1) at a ratio of 25:1 (MBP-Smt3-*Ml*GH17A /Ulp1 (w/w)) for 1 h at 30 °C. The protein was then purified by metal ion affinity chromatography using a HisTrap HP 1ml column (GE Healthcare Life Sciences), with 20 mM Tris–HCl, 10 mM imidazole, pH 7.4, 500 mM NaCl as binding buffer, and 20 mM Tris–HCl, 500 mM imidazole, pH 7.4, 500 mM NaCl as elution buffer. The protein was eluted by applying an imidazole gradient up to 500 mM over 10 column volumes. Fractions containing the recombinant protein were immediately collected and stored at 4 °C. The protein concentration was determined considering the absorption coefficient (91790 M^-1^Cm^-1^) (30) by measuring A_280_ using a NanoDrop spectrophotometer (Thermo Scientific). The integrity and purity of the protein were analyzed by 4–15% glycine-SDS-PAGE. Fractions containing the pure protein were combined and kept at 4 °C for further analysis.

### Thermal unfolding temperature determination

The protein’s thermal unfolding transition temperature (T_m_) was determined using nano-differential scanning fluorimetry (nanoDSF) on a Prometheus NT.48 instrument. The enzyme was diluted to a concentration of 0.2 mg/mL in 20 mM phosphate buffer, pH 7.4, with 200 mM NaCl before the analysis. The fluorescence intensity ratio of 350/330 nm was continuously monitored at 30% intensity while subjecting the sample to a temperature gradient of 1.0 °C/min, ranging from 20 °C to 95 °C. The T_m_ was determined by identifying the inflection point on the unfolding curve. Furthermore, the protein mid-aggregation temperature (T_onset_) was measured by estimating light scattering. This experiment was performed in triplicate, and the average values with the corresponding standard deviation were reported.

### Temperature and pH optima

The enzyme’s optimal condition was determined by measuring the activity in a pH range of 3.6-6 using laminarin as the substrate. Reaction mixtures of 60 μL were prepared by mixing 6 μL of laminarin 20 mg/mL, 6 μL of 500 mM buffer (acetate buffer for pH 3.6–5.5 and potassium phosphate buffer for pH 6.0), 24 μL of the enzyme, and 24 μL of ultrapure water (Milli-Q grade). The reaction mixtures were then incubated in a PCR machine (SensoQuest), and subjected to a temperature gradient ranging from 13 °C to 53 °C, for 30 min. To stop the reaction, 120 μL of 3,5-Dinitrosalicylic acid (DNS) was added, and the reaction mixture was incubated at 95 °C for 10 min. After cooling, the absorbance at 540 nm was measured using a microplate spectrophotometer (Thermo Fisher Scientific) to determine the amount of reducing ends, with glucose serving as the standard. Duplicate experiments were performed for each condition.

### Product profile analysis

The amount of oligosaccharides obtained from the enzymatic reaction of *Ml*GH17A was analyzed by employing a high-performance anion exchange chromatography with pulsed amperometric detection (HPAEC–PAD) system (Thermo Fischer Scientific) coupled with a PA-200 column and pre-columns. Oligosaccharides were separated using eluents as follows: (A) Ultrapure water (Milli-Q grade), (B) 1M NaOAc in 200 mM NaOH, and (C) 200 mM NaOH. The separation was obtained using a linear gradient of (B) up to 500 mM NaOAc in (C), while the eluent (A) was maintained at 50% over 25 min; subsequently, the eluent (B) was increased to 100% for 5 min. The flow rate was set to 0.5 mL/min, and the column compartment was kept at 30 °C during the analysis. To construct a calibration curve, laminari-oligosaccharides were utilized as standards.

The enzyme mode of action (endo or exo activity) was assessed by performing the enzymatic reaction with a very low concentration of the enzyme using laminarin as the substrate. Samples were taken at very short intervals (10-30 seconds) and the reaction products were analysed using HPAEC-PAD.

### Enzyme kinetic determination

The kinetic parameters were determined using laminarin as a substrate. Following the procedure described in the “Temperature and pH optima” section, reaction mixtures of 60 μL were prepared using 500 mM acetate buffer pH 4.5. Fourteen different concentrations of substrate, ranging from 0 to 4 mM, were selected, and the final concentration of the enzyme in the reaction mixtures was set at 1 nM. The reactions were conducted at 40 °C for 10 min, followed by boiling at 95 °C for 5 min to terminate the reaction. To calculate the kinetic parameters, 120 μL of DNS (3,5-Dinitrosalicylic acid) was added to the reaction mixtures, followed by boiling at 95 °C for 10 min. Standard samples were prepared by mixing 6 μL of 500 mM acetate buffer pH 4.5, 6 μL of glucose (ranging from 0 to 5 mM), and 24 μL of water; after adding 120 μL of DNS, 24 μL of the enzyme (at a final concentration of 1 nM) was added, and the standard samples underwent the same boiling procedure as the reaction mixtures. The amount of reducing end was determined by measuring the absorbance at 540 nm. Kinetic parameters were determined by a non-linear regression according to the Michaelis Menten model using GraphPad Prism software version 9.4.0 for Windows (GraphPad Software, San Diego, California, USA, www.graphpad.com).

### Nuclear Magnetic Resonance Spectroscopy

1D and 2D NMR spectra were recorded on a 600 MHz Bruker Avance Neo NMR spectrometer equipped with a 5mm TCI Prodigy CryoProbe (Bruker) at 311K. Samples were dissolved in D_2_O/PBS buffer (0.5 mL 99.9%; Cambridge Isotope Laboratories, Tewksbury, MA, USA) and transferred to 5 mm NMR tubes. All spectra were obtained by using standard Bruker pulse programs with water suppression. The 2D TOCSY spectra were recorded using an MLEV-17 mixing sequence with spin-lock times of 150 ms. The 2D NOESY spectra were recorded with a mixing time of 300 ms. Chemical shifts were expressed in parts per million (ppm) relative to internal acetone (δ 2.225 for ^1^H and δ 31.07 for ^13^C). Data were processed by using TopSpin^TM^ software (Bruker). The 2D tr-NOESY spectra were acquired in a phase-sensitive mode using the States-TPPI method and additional water signal suppression by presaturation. Time domain data were collected with 256 increments (t1 dimension) and 2048 datapoints during the acquisition period with a mixing time of 50 ms - 400 ms and relaxation delay of 2 s. The optimal mixing time of the tr-NOESY experiment was set to 200 ms.

Two tr-NOESY spectra were acquired, one with octasaccharide-branched ligand in the presence of the *Ml*GH17A receptor and the other in the absence of *Ml*GH17A receptor. After optimization of ligand/protein ratio, the final sample was prepared with 0.03 mM protein and 3 mM of the octasaccharide-braanched product (100:1 ligand/protein ratio) in 500 mL of phosphate buffered saline (pH 5.0).

### Matrix-Assisted Laser Desorption/Ionization Time-of-Flight Mass Spectrometry (MALDI-TOF/TOF MS)

MALDI-TOF/TOF MS experiments were performed using Bruker ultrafleXtreme (Bruker Daltonics) mass spectrometer. All spectra were recorded in reflector positive-ion mode and the acquisition mass range was 200–6000 Da. Samples were prepared by mixing on the target 0.5 μL sample solutions with 0.5 μL aqueous 10% 2.5-dihydroxybenzoic acid as matrix solution.

Tr-NOESY NMR Experiments—To get more information on the bound conformation of the octasacharide-branched ligand with *Ml*GH17A, tr-NOESY experiments were recorded in the ligand/protein ratio 100:1.

The tr-NOESY experiment showed a difference of the NOE cross-peak patterns between the free and bound states. The large octasaccharide-branched ligand in free state, have correlation times of several nanoseconds and, therefore, display negative NOEs at frequencies of 600 MHz.

On the other hand, when ligand is bound to a protein receptor, gives positive NOEs at δ 3.37 (H-2 of terminal Gt residue), at δ 3.52 and δ 3.51 (H-2 and H-4, respectively, of internal GC residue) and at δ 3.89 (H-6b branched GD residue), suggesting close proximity of these protons to catalytic active site of *Ml*GH17A.

The complete assignment of chemical shifts of the protons of the free octasaccharide-branched ligand was previously described by Allahgholi et al. (10).

### Bioinformatics analysis

#### Sequence alignment

The complete sequence of the protein was used for the sequence alignment. The amino acid sequences of the laminarinases from the GH17 family, including *Fa*GH17A (GenBank CDF79584.1), from *Formosa agariphila*; *Fb*GH17A (GenBank AOR29489.1), from *Formosa* sp. Hel1_33_131; *Fb*GH17B (GenBank AOR29491.1), from *Formosa* sp. Hel1_33_131; and *Vb*GH17A (GeneBank OEF87991.1), from *Vibrio breoganii*; *Vv*GH17, β-1,3-glucanase from *Vibrio vulnificus* (GenBank ASM98089.1); and the β-1,3-glucanosyltransglycosylase (*Ml*GH17B) from *M. lutaonensis* ISCAR-4703 (GenBank WIW39500) were obtained, and the alignment was performed using Clustal Omega (https://www.ebi.ac.uk/Tools/msa/clustalo/) through Jalview version 2.11.2.6 using the ClustalO web service with default settings (31).

#### Homology modeling

The three-dimensional structure of *Ml*GH17A was modeled by homology modeling using the YASARA program (YASARA Biosciences) as described in Linares-Pastén et al. (21). The crystal structure of a laminarinase from *Formosa* sp. Hel1_33_131 *(Fb*GH17A) (PDB 6FCG), which has 64 % sequence identity to *Ml*GH17A, was used as the template, and modeling parameters were set according to supplementary table S4.

Model refinement was carried out for the generated hybrid model using default parameters in the YASARA program. A simulation cell was created with dimensions 2×7.5 Å larger than the model along each axis and was filled with water molecules, and Na^+^ and Cl^-^ ions were included as counter ions. The simulation was conducted for 500 ps using the YAMBER03 force-field, and snapshots were saved at intervals of 25 ps.

The quality of the refined structures was evaluated by programs available at UCLA-DOE LAB— SAVES v6.0 (saves.mbi.ucla.edu/). The average 3D-1D score was calculated using Verify 3D (32, 33). Ramachandran plot was generated using PROCHECK (34). Overall quality factors were also assessed using ERRAT (35).

The three-dimensional model of the enzyme was visualized and presented using the PyMOL v2.5.2 program developed by Schrödinger.

#### Docking of laminari-oligosaccharides

Docking studies were conducted using Glc_5_ as the shortest substrate, considering that the activity on Glc_4_ was very low.

A β-1,3 linked oligosaccharide with DP5 was built using the oligosaccharides building tool available in the YASARA program. After MD simulation at physiological conditions (pH 7.4 and 298 K) for 2 ns, the conformer with the lowest forcefield energy was selected for docking studies. Docking of the enzyme with the designed ligand was conducted with the docking locally implemented in YASARA. The docked structure was subjected to the model refinement using default parameters in YASARA and the model with the highest score was selected for further analysis. Furthermore, the ligand was manually extended by adding glucose moieties to putative subsites at both the reducing and non-reducing ends of the ligand, in a proper conformation via a β-1,3 linkage, for the investigation of the enzyme’s putative subsites. The ligand was hydrolyzed manually and subjected to energy minimization. The enzyme ligand interactions were studied by Chimera version 1.17.1 (36) and Ligplot^+^ (37).

The three-dimensional model of the enzyme-ligand complex and the active site were graphically presented using the PyMOL v2.5.4 program (Schrödinger).

#### Protein subcellular location prediction

The presence of signal peptidase I and II cleavage sites was investigated using SignalP 5.0 (services.healthtech.dtu.dk/services/SignalP-5.0/) (38) and LipoP 1.0 (services.healthtech.dtu.dk/services/LipoP-1.0/) (39) servers, respectively. The subcellular location of the protein was investigated using CELLO (subCELlular LOcalization predictor) v.2.5 (40) and PSORTb v.3.0.

## Supporting information

Additional supplementary figures, and tables are provided in the supporting information.

## Acknowledgment

Funding was obtained from the ERA-NET Cofund BlueBio MariKat project, from the Swedish Research Council Formas (grant No. 2019-02359) and from the SBEP FunSea (grant No 2023-02261, as well as from the European Union’s Horizon Europe program SeaMark project (grant No. 101060379). Authors would like to express gratitude to Elísabet Eik Guðmundsdóttir, Edda Olgudóttir and Léonie Jouy for technical assistance.

## Author Contributions

E.N.K., L.A. conceptualization; L.A., A.M., M.G.N.D., J.M.D., Z.D., J.A.L.-P., and E.N.K. formal analysis; E.N.K., and G.O.H. funding acquisition; L.A., investigation; L.A., A.M. and J.M.D. methodology; L.A. data curation, validation and visualization; E.N.K, G.O.H. project administration, E.N.K, O.H.F. resources; E.N.K, O.H.F supervision; L.A. writing original draft; L.A., M.G.N.D., J.M.D, Z.D., J.A.L.-P., O.H.F., A.M., G.O.H and E.N.K. writing, review, and editing.

